# Cell-type specific effects of *Fusarium* mycotoxins on primary neuronal and astroglial cells

**DOI:** 10.1101/2024.02.29.582764

**Authors:** Viktória Szentgyörgyi, Norbert Bencsik, Brigitta Micska, Anikó Rátkai, Katalin Schlett, Krisztián Tárnok

## Abstract

Fumonisin B1, deoxynivalenol (DON) and zearalenone (ZEA) are toxic secondary metabolites produced by *Fusarium* molds. These mycotoxins are common food and feed pollutants and represent a risk for human and animal health. Although the mycotoxins produced by this genus can cross the blood-brain-barrier (BBB) in many species, their effect on neuronal function remains unclear. We investigated cell viability effects of these toxins on specified neural cell types, including mouse primary neuronal, astroglial and mixed cell cultures 24 or 48 hours after mycotoxin administration. Cell viability assay revealed that DON decreased cell viability in a dose-dependent manner, independently from the culture’s type. Fumonisin B1 increased cell viability significantly on astroglial and mixed cell cultures in lower doses, while it exerted a highly toxic effect in 50 µM. ZEA had significant effects on all culture type in 10 nM by increasing the cell viability. Since ZEA is a mycoestrogen, we analyzed the effects of ZEA on the expression of estrogen receptor isotypes ERα and ERβ and mitochondrial voltage-dependent anion channel (VDAC1) by qRT-PCR. In neuronal and mixed cultures, ZEA administration decreased ERα expression, while in astroglial cultures, it induced the opposite effect. ERβ and VDAC1 expression was not altered by ZEA in either culture types. ZEA also affected the firing pattern of neurons by enhancing the burst frequency. Our results demonstrate that *Fusarium* mycotoxins are acting on a cell specific manner in the brain tissue.

## Introduction

Fumonisin B1 (FB1), deoxynivalenol (DON) and zearalenone (ZEA) mycotoxins are common pollutants in food and feed worldwide and represent risks for both human and animal health (Marin et al. 2013). These toxic secondary metabolites are produced by molds of the *Fusarium* genus. Due to climate change, the habitat of these species alters, and the new weather conditions supports their mycotoxin production (Farkas et al. 2011). Mycotoxins are usually thermostable, therefore, they can be present in the processed food and feed despite of milling, extrusion, storage or heating. Mycotoxins can induce a multitude of physiological effects. For example, FB1, the most prevalent fumonisin-type, is a known cancer promoter (Gelderblom et al. 2001) linked to human esophageal cancer in South Africa and China. It is hepato-, nephro- and immunotoxic as well, caused by the mechanisms of oxidative stress, apoptosis and inhibition of the sphingolipid biosynthesis (Stockmann-Juvala and Savolainen 2008). Chronic exposure to DON leads to feed refusal, anorexia, immunotoxicity, gastroenteritis and skin dermatitis based on its inhibitory action on protein biosynthesis and sub-sequent activation of mitogen-activated protein kinases (Pestka 2007, 2010). ZEA also has immuno- and hepatotoxic effects and induces genotoxicity by DNA-adduct formation, as well (Zinedine et al. 2007). ZEA is a potent endocrine disruptor due to its binding capacity to both the ERα and ERβ estrogen receptors (Takemura et al. 2007; Ropejko and Twarużek 2021). By misregulating the hormonal system, it can cause severe symptoms such as fertility disorders, reduced litter size and fetal weight, alteration in the reproductive track in animals or early thelarche and premature puberty in humans (Zinedine et al. 2007).

Besides the general physiological effects, all of these mycotoxins can cause neurological symptoms (Marin et al. 2013). In the case of FB1, the best-known effect is the equine leukoencephalomalacia (ELEM) in horses (Domijan 2012), however, FB1 has also been linked to neural tube defects in children (Missmer et al. 2006). Former studies also showed that FB1 can pass through the blood-brain-barrier (BBB) (Domijan 2012). *In vitro* studies on neural cultures demonstrated that FB1 causes a large scale of cellular effects. It can inhibit axonal and dendritic growth (Harel and Futerman 1993; Furuya et al. 1995) or alter myelin formation and oligodendrocyte maturation (Monnet-Tschudi et al. 1999) by disrupting sphingolipid metabolism (Soriano et al. 2005). Based on data from different cell lines, FB1 can induce cell death by oxidative stress and/or apoptosis in a cell type specific manner (Doi and Uetsuka 2011).

Due to its strong emetic properties, deoxynivalenol is also known as vomitoxin. It is able to exert a direct neuronal effect, since circulating DON can pass the BBB and accumulates in the brain (Pestka et al. 2008; Maresca 2013). DON has been reported to cause GABAergic neuronal activation (Csikós et al. 2020) and stimulate the dopamin receptors of neurons (Prelusky et al. 1992; Sobrova et al. 2010). *In vivo*, it has also been reported to increase salivation in swine (Rotter et al. 1996). DON can cause neuroinflammation (Girardet et al. 2011a), can modify feeding behavior leading to reduced food intake (Collins et al. 2006; Girardet et al. 2011b) and can alter goal-directed, reward-driven behaviors in rats (Csikós et al. 2020).

Few data have been reported about direct neurologic and/or neurotoxic effects of ZEA. *In vivo*, chronic ZEA administration decreased the brain protein levels of rats and affected the oxidoreductive capacity of cells by increasing superoxide dismutase (SOD) and decreasing glutathione (GSH) levels and glutathione peroxidase activity (GSH-Px) resulting in elevated level of NO and apoptotic cells (Ren et al. 2016). ZEA-induced formation of reactive oxygen species (ROS) and oxidative stress were also observed in human neuroblastoma cell lines (Venkataramana et al. 2014; Agahi et al. 2020b, a). On the other hand, ZEA exerted no effect on estrogen biosynthesis in glioblastoma cells (Wang et al. 2014). In a few reports, administration of ZEA or its derivatives led to positive outcome on memory impairment and adult neurogenesis (Dong et al. 2014) or was neuroprotective by counterbalancing the effects of ovariectomy or β-amyloid (25-35) induced oxidative / ER stress (Dong et al. 2006, 2007, 2017). Regulation of the expression of ERβ or thyroid receptor (α,β) by ZEA has also been recently reported on cerebellar granule cells (Jocsak et al. 2016; Kiss et al. 2018). These data reveal important neurological consequences of mycotoxin consumption but we have insufficient information on cell-type specific effects within the brain.

Purified mycotoxins are intensively tested on different cell lines to determine their primary effects on cell survival and metabolism (Juan-García et al. 2013; Smith et al. 2018). On the other hand, primary cultures enable to investigate cells derived directly from the brain tissue. To unravel the neuron or astroglia specific processes upon mycotoxin-treatment, pure neuronal and astroglia cultures as well as mixed cultures containing neurons and astroglial cells were established and analyzed. In the present work, we compare the cell-type specific effects of FB1, DON and ZEA to define cellular targets of the different mycotoxins. We provide evidence that neurons and astrocytes can counterbalance mycotoxin-evoked effects on each other. Additionally, we show that ZEA promotes spontaneous bursting activity of neuronal networks.

## Materials and methods

### Animal handling

CD1 wild-type mice were purchased from Charles River Laboratories (Wilmington MA, USA; organism: RRID:IMSR_CRL:22). Mice were housed at 22 ± 1°C, with 12 h light/dark cycles and *ad libitum* access to food and water. All experiments complied with local guidelines and regulations for the use of experimental animals (PEI/001/1108-4/2013 and PEI/001/1109-4/2013), in agreement with EU and local legislation.

### Drug preparation

Fumonisin B1 (FB1, Tocris), deoxynivalenol (DON, Tocris) and zearalenone (ZEA, Tocris) were dissolved in 10% DMSO (FB1), 10% ethanol (DON) or in 10% ethanol-50% DMSO (ZEA), respectively. Cells were treated with the mycotoxins in 1 nM-50 μM concentration range, for 24 or 48 hours. Control wells were treated with the correspondent solvent.

### Preparation of cell cultures

The experiments are performed on mixed (where both neurons and astroglia cell were present) and on pure neuronal or astroglial primary cultures. Primary cultures of embryonic cortical neurons were prepared from pregnant CD1 mice on embryonic days 14–15, according to Tárnok et al. (2008). Briefly, under aseptic conditions, cortices were isolated, cleaned from meninges and were incubated in 0.05% trypsin-EDTA solution (Gibco, Thermo Scientific) for 15 min at 37°C. After a brief centrifugation step, cells were triturated in NeuroBasal media (Gibco, Thermo Scientific) supplemented with 2% B27 (Gibco, Thermo Fisher), 5% FCS (PAN-Biotech, Germany), 0.5 mM Glutamax (Gibco, Thermo Scientific), 40 μg/ml gentamycin (Hungaropharma, Hungary) and 2.5 μg/ml amphotericin B (Sigma, Hungary), and filtered through a sterile polyester mesh with 42 μm pore size (EmTek Ltd, Hungary). Cells were seeded onto poly-L-lysine (PLL; Sigma, Hungary) coated 96 well plates (Greiner, Hungary) at 8×10^4^ cells/well density. For microscopy, 1.5×10^5^ cortical neurons were seeded onto poly-L-lysine – laminin (1 μg/cm^2^ Sigma, Hungary) coated glass coverslips in 24-well plates. Cells were cultivated at 37°C in 5% CO_2_/95% air atmosphere. Neurons were cultured in Neurobasal medium (Life Technologies, Hungary), supplemented with 2% B27 (Life Technologies, Hungary), 0.5 mM Glutamax (Gibco, Thermo Scientific), 40 μg/mL gentamycin (Hungaropharma, Hungary), and 2.5 μg/mL amphotericin B (Sigma, Hungary). 5% FCS (PAN-Biotech, Germany) was included in the culture medium until DIV1 (pure neuronal cultures) or DIV4 (mixed neuronal-glial cultures). Complete and partial (1:2) medium changes were performed on DIV1 and DIV4, respectively. Pure neuronal cultures were treated with cytosin- arabinofuranoside (CAR, 10 μM; Sigma, Hungary) 24 h after plating to prevent the further division of nonneuronal cells. In the case of mixed cultures, cells were treated with 10 μM CAR only on the 6^th^ day after plating. Mycotoxin-treatments were performed on the 7-8^th^ day of cultivation (details in the text).

Primary cultures of astrocytes were prepared from 1–4 days old CD1 mice, according to Tárnok et al. (2010). Briefly, cerebral cortices were isolated, freed from meninges and incubated in trypsin-EDTA [0.5 mg/mL in phosphate-buffered saline (PBS), Gibco, Thermo Scientific, Hungary] and 5 mg/mL DNase solution (Sigma, Hungary) for 15 min at 37°C. After a brief centrifugation, cells were triturated in high glucose Dulbecco’s Modified Eagle Medium (HDMEM, Sigma, Hungary) supplemented with 10% fetal calf serum (FCS, PAN-Biotech, Germany), 40 μg/mL gentamycin, 2.5 μg/mL amphotericin B (all from Sigma, Hungary) and filtered through a sterile polyester mesh with 42 μm pore size (EmTek Ltd, Hungary). Cell number was determined by trypan blue exclusion. Cells were seeded onto poly-L-lysine (Sigma) coated Petri dishes at 3–4×10^5^ cells/cm^2^ density in DMEM containing 10% FCS, 2 mM L-glutamine, 2.5 μg/mL amphotericin B and 40 μg/mL gentamycin. Cells were cultivated at 37°C in 5% CO_2_/95% air atmosphere. Astroglial cultures were used for further experiments after reaching confluency (7–8 days *in vitro*). For the experiments, cells were reseeded into poly-L-lysine-coated 96-well plates or onto 12mm diameter coverslips in 24-well plates in 2.5×10^4^ cells/cm^2^ density.

### Cell viability assay

Viability of the cultures was measured 24 h or 48h after mycotoxin treatment by MTT assay as described by Tárnok et al. (2008), Briefly, cells grown in 96-well plates were treated with 3-(4,5-dimethylthiazol-2-yl)-2,5-diphenyltetrazolium bromide (MTT, Sigma, Hungary) in a final concentration of 200 μg/ml. After 35-40 min incubation, cells and formazan crystals were dissolved in acidic (0.08M HCl) isopropanol (Merck, Hungary). Optical density was determined at a measuring wavelength of 570 nm against 630 nm as reference with a Multiscan EX ELISA reader (Thermo, Bio-science, Hungary). Assays were carried out at least on 6 parallel wells of 3 independent cultures. Viability data were determined as averages and standard errors of the mean and were expressed as a percentage of the control cultures. Data were compared by Student’s t-test (p<0.05).

### Immunocytochemistry and microscopic analysis

Cells grown on 12 mm diameter coverslips in 24-well plates were fixed with 4% paraformaldehyde (Taab, UK; w/v in PBS), permeabilized with 0,1% Triton-x-100 in phosphate-buffered saline and blocked in 2% bovine serum albumin in phosphate-buffered saline.

To detect the different cell-types in neural cultures, cells were immunostained with the anti-IIIβ-tubulin (mouse, 1:1000; EXBIO, Czech Republic), anti-GFAP (chicken, 1:1000, Synaptic System, Germany) and anti-Iba1 (goat, 1:500, Novusbio, Bio-Techne, Hungary) antibodies overnight at 4°C. Anti-mouse Alexa Fluor-DyLight594 (Vector Laboratories, UK), anti-chicken Alexa Fluor-647 (Invitrogen, Thermo Scientific) and anti-goat Alexa Fluor-488 (Invitrogen, Thermo Scientific) were applied in a 1:500 dilution for 1h at RT. Preparations were mounted with bis-benzimide (Sigma, Hungary)-containing Mowiol 4.88 (for nuclei labeling, Polysciences, Germany).

To stain mitochondria of astroglia cells, 10 nM MitoTracker Orange (Thermo Scientific, USA) was applied at 37°C for 25 min before fixing (in the presence or absence of ZEA). After the fixation, cell membranes were labeled by CholeratoxinB-Alexa647 (50 μg/mL, Invitrogen, Thermo Scientific) for 30 min at RT. Preparations were mounted with bis-benzimide - containing Mowiol 4.88 (for nuclei labeling, Polysciences, Germany).

Microscopic images were acquired with an inverted AxioObserver Z1 wide-field fluorescent microscope (Zeiss, Hungary) with a LD Plan Neofluar 20×/0.4 NA (Zeiss, Hungary) or an α-PlanApoChromat 63×/1.46 NA oil immersion objective (Zeiss Hungary) using Colibri-LED illumination and the appropriate filter sets. Images from sister cultures for the same experiment were recorded with identical microscope settings and analyzed in the same way.

Mitochondrial area was determined in projected z-stack images of astrocytes taken by 63× magnification. CholeratoxinB and DAPI staining was used to delineate cell shape and nucleus, respectively using Fiji software (Schindelin et al. 2012). MitoTracker Orange positive objects within the cytoplasm were separated from the background by intensity thresholding and their total area was determined to calculate the total mitochondrial area / cytoplasmic area ratio.

### Quantitative real-time PCR (qRT-PCR)

Cultures were lysed and RNA samples were obtained using Quick-RNA MiniPrep (ZYMO Research, USA). Reverse transcription was performed with the Maxima First Strand cDNA Synthesis for RT-qPCR kit (Thermo Scientific) according to the manufacturer’s instruction. Messenger RNA expression was investigated using the Maxima SYBR qPCR Master Mix (Thermo Scientific) with specifc primers (see Supp. Table 1.). The qPCR run was performed using a CFX96 (C1000 Touch) from Bio-Rad Laboratories (USA) with the following settings: 1 cycle at 95°C for 10min, 40 cycles at 95°C for 15s followed by 55°C for 30 s and 72°C for 30 sec. Cq and RFU values were obtained from Bio-Rad CFX Manager software (Bio-Rad, UDSA). The relative expression of the interested genes was calculated using the ΔΔCq method using triplicated samples.

### MEA recordings and analysis

In MEA experiments, cells were seeded onto MEA60-4Well-PT (Qwane Biosciences, Switzerland), 60ThinMEA 100/10 ITO (Multichannel Systems, Germany) or 60MEA 200/30 ITO (Multichannel Systems, Germany) multi-electrode chambers at 4×10^4^ (MEA60-4Well-PT) or 4-5×10^5^ (60ThinMEA 100/10 ITO and 60MEA 200/30 ITO) cells/well densities. Prior to seeding the cells, surface of the multi-electrode arrays was coated by poly-L-lysine (2 µg/cm^2^, Sigma, Hungary) and laminin (4.2 µg/cm^2^, Sigma, Hungary).

Spontaneous extracellular field potentials were recorded at 37 °C (Temperature Controller TC01102, Multichannel Systems) using a USB-1060 INV Microelectrode array System (Multichannel Systems, Germany) with a MEA Amplifier with Blanking Circuit (Multichannel Systems, Germany) amplifier at a sampling rate of 20 kHz/channel. Signals were high-pass filtered at 300 Hz and low-pass filtered at 3000 Hz with a Butterworth 2^nd^ order filter. Spontaneous firing was recorded for 10 minutes before the addition of the toxins (baseline) and repeated after the toxin treatment for another 10 minutes. Recorded spikes were detected by a threshold crossing method (Thr.%: −1.1) and were sorted by the *k means* algorithm. The Thr.% shows the threshold value in A/D counts and the associated voltage that was used to collect the waveforms. The spike timestamps were extracted using the Plexon Offline Sorter (Plexon Inc., Texas, USA). Analysis of the spike timestamps, including characterizing spontaneous firing and burst activity was performed by the NeuroExpress software (https://www.researchgate.net/project/Analysis-software-for-whole-cell-electrophysiological-data).

### Statistics

For statistical evaluation of pairs of data sets, Student’s t-tests (paired or independent, in case of normal distribution of data) or non-parametric Mann–Whitney tests (in case of non-normal distribution of data) were used. Statistics were calculated using SPSS Statistics (IBM). A p-value equal to or smaller than 0.05 was considered statistically significant. Number of independent experiments (n) is indicated in the figure legends.

## Results

### The presence of astroglial cells attenuates the cytotoxic effect of FB1

Neurons and astrocytes can exhibit diverse responses to mycotoxin administration. In order to determine and compare cell-type specific effects of mycotoxins, three different types of primary cell cultures were created and used in our experiments containing either i) both neurons and astroglia (“mixed”), ii) only neurons and no glial cells (“neuronal culture”) or iii) neuron-free glial cells (“astroglial culture”). The cellular composition of the cultures was verified by immunostaining using cell type specific markers. Neurons, astroglia cells and microglia were visualized by IIIβ-tubulin, glial fibrillary acidic protein (GFAP) and Iba-1 immunopositivity, respectively (Fig. 1). In mixed cultures, both neurons and astrocytes were detectable. Neuronal cultures were free of astroglial and microglial cells and while primary glia cultures contained a few microglial cells, they were mainly astrocyte-rich and neuron-free.

**Figure 1.**
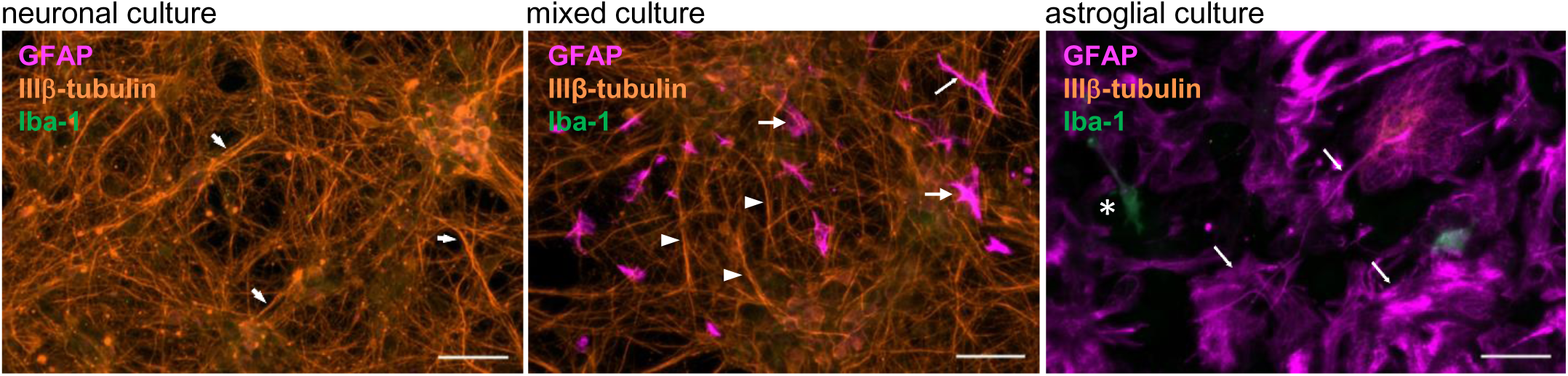
The representative immunofluorescence images of the neural cultures applied for the mycotoxin measurements. Neurons (arrowheads), astroglial cells (arrows) and microglia (asterisk) were visualized by IIIβ-tubulin, glial fibrillary acidic protein (GFAP) and Iba-1 immunopositivity, respectively. Neuronal cultures were free of astroglial and microglial cells on DIV7 since mitosis was inhibited 24 hours after the plating by adding CAR. In mixed cultures, both neurons and astrocytes were detectable on DIV7. Astroglial cell cultures contained less than 2% microglia (asterisk) and they were free of neurons 24 hours after seeding. Scale bars, 50 μm.

To examine cellular physiological effects of FB1 mycotoxin, mixed and purified cultures in 96-well plates were treated for 24 or 48 hours with increasing (1 nM-50 µM) concentrations of FB1 and cell viability was determined by MTT assay. According to our data, FB1 significantly increased cell viability above 10 nM concentration in cultures containing solely neurons or neurons and astroglia cells mixed (Fig. 2A, B). However, from 10 µM concentration, this effect was decreased and in pure neuronal cultures, 50 µM FB1 caused a drastic, 50% viability-loss. Increasing the treatment from 24 to 48 hours did not change dramatically the observed effects of the toxin.

**Figure 2.**
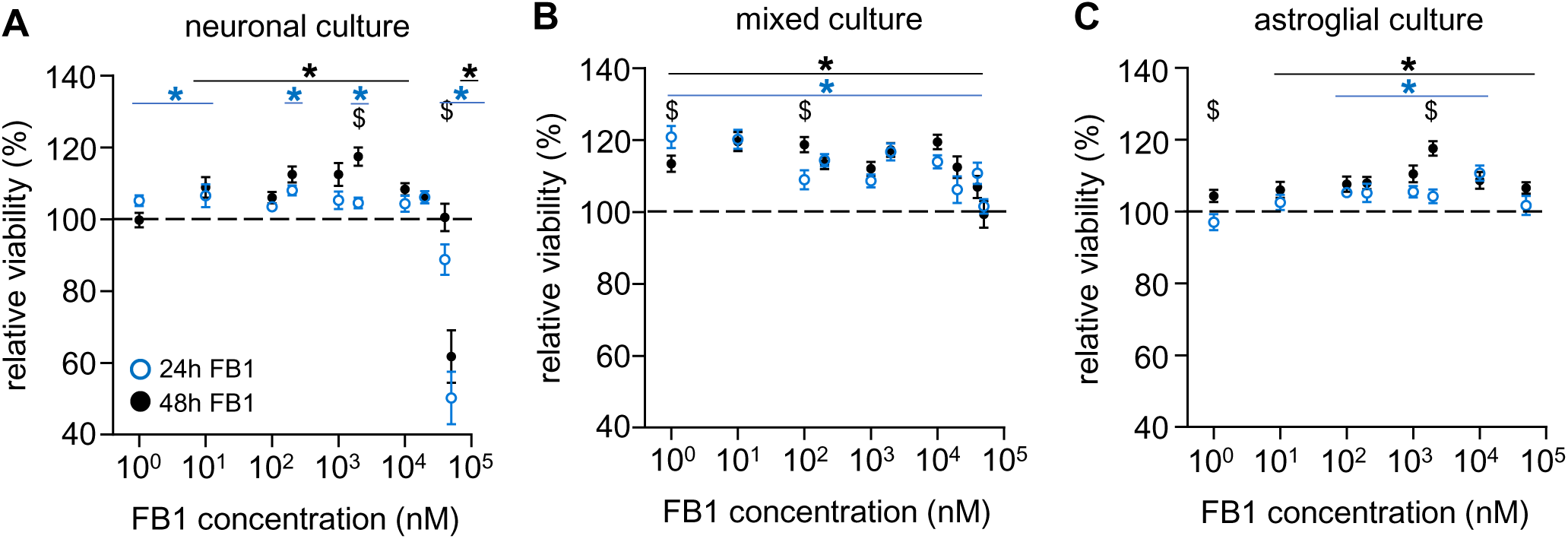
Astroglial cells attenuate the cytotoxic effect of FB1 observed in neurons. The effects of FB1 on the cell viability of mouse primary (A) neuronal, (B) mixed and (C) astroglial cell cultures were determined after 24- or 48-hours treatments of 1, 10, 100, 200, 1000, 2000, 10000 or 50000 nM FB1 by MTT assay. Results are expressed in the percentage of control data, as mean ± s.e.m. „$” indicates significant differences between data obtained after 24- and 48-hours treatment; * indicates significant differences compared to non-treated data ($;*:p<0.05). n=12-42.

In astroglial cultures, FB1-administration significantly increased cell viability above 10 nM or 100 nM concentration in 48 or 24-hours treatment-time, respectively (Fig. 2C). Cytotoxic effects observed in pure neuronal cultures did not occur in glial nor in mixed cultures, so the presence of astroglial cells reduced the neural cytotoxicity of FB1.

### Astroglial cells are more sensitive than neurons to the cytotoxic effects of DON

In addition to FB1, *Fusarium* species also produce other mycotoxins including deoxynivalenol (DON). To investigate the effects of DON on neural cells, cultures were treated similarly to that used in the FB1 experiments. Our results showed that in cultures containing neurons, DON treatment at concentrations below 1 µM was ineffective (Fig. 3A, B). In the concentration range of 1-50 µM, DON decreased cell viability in a concentration- and time-dependent manner. On the other hand, a more prominent, concentration-dependent decrease in viability was observed in astroglia cells already after 24 hours of DON treatment above 200 nM (Fig. 3C). Toxicity was further elevated by 48 hours of DON treatment and was evident from even lower concentrations (e.g., at 2 µM, 48 hours treatment of DON resulted in ~ 35% reduction in viability of astroglia cells) (Fig. 3C). Notably, in mixed cultures, DON treatment reduced cell viability only by 20% at 2 µM (Fig. 3B). Taken together, our data show that astroglial cells are the most sensitive cell to DON administration in our system.

**Figure 3.**
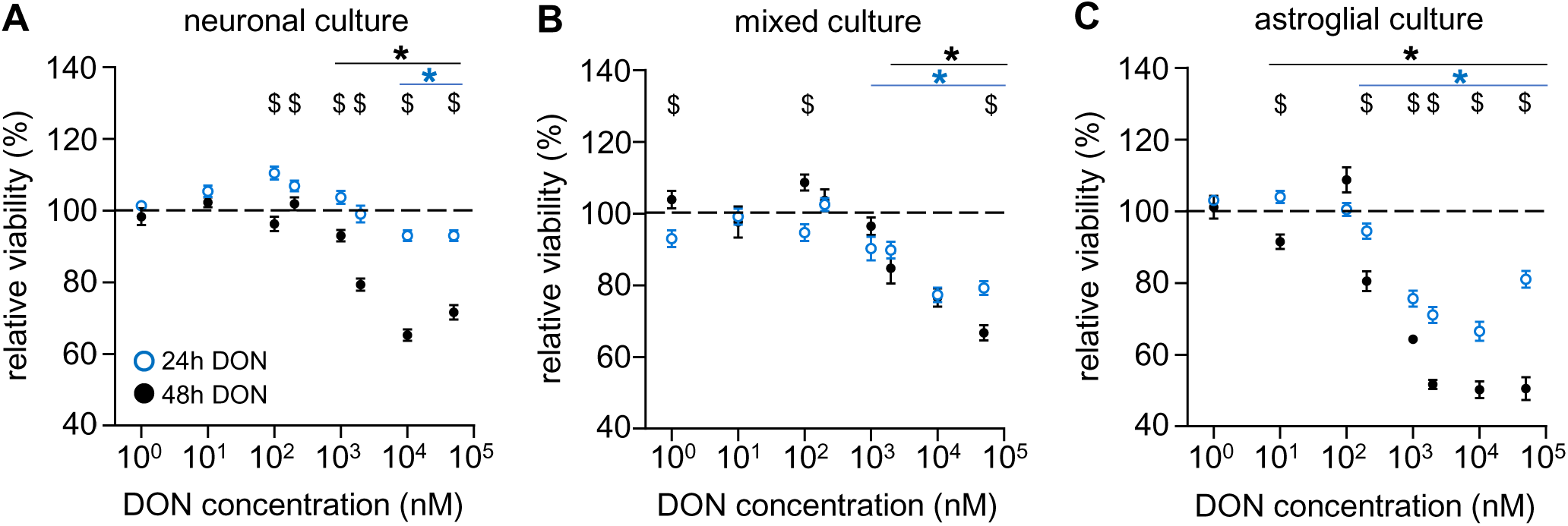
Astroglial cells are the most sensitive to the cytotoxic effect of DON. The effects of DON on the cell viability of mouse primary (A) neuronal, (B) mixed and (C) astroglial cell cultures were determined after 24- or 48-hours treatments of 1, 10, 100, 200, 1000, 2000, 10000 or 50000 nM DON by MTT assay. Results are expressed in the percentage of control data, as mean ± s.e.m. „$” indicates significant differences between data obtained after 24- and 48-hours treatment; * indicates significant differences compared to non-treated data ($;*:p<0.05). n=18-36.

### ZEA increases cell viability in all investigated culture types even at low concentrations and alters the expression of ERα in a cell type-specific manner

The physiologically relevant concentration of ZEA in brain tissue is predicted to be in the 1-10 nM range in rodents, feeding the animals with no-observed-effect level (NOEL) doses (Shin et al. 2009b). Former studies also showed that ZEA is bioactive even at low, 10 nM concentration in several cell-based system (Phelps et al. 1988; Dong et al. 2007; Zheng et al. 2007, 2018). This observation was confirmed by our preliminary screening (data not shown), so we detailed analysis were performed using 10nM concentration. According to our data, 48 hours treatment with ZEA at 10 nM significantly increased the viability of each culture type (Fig. 4A).

**Figure 4.**
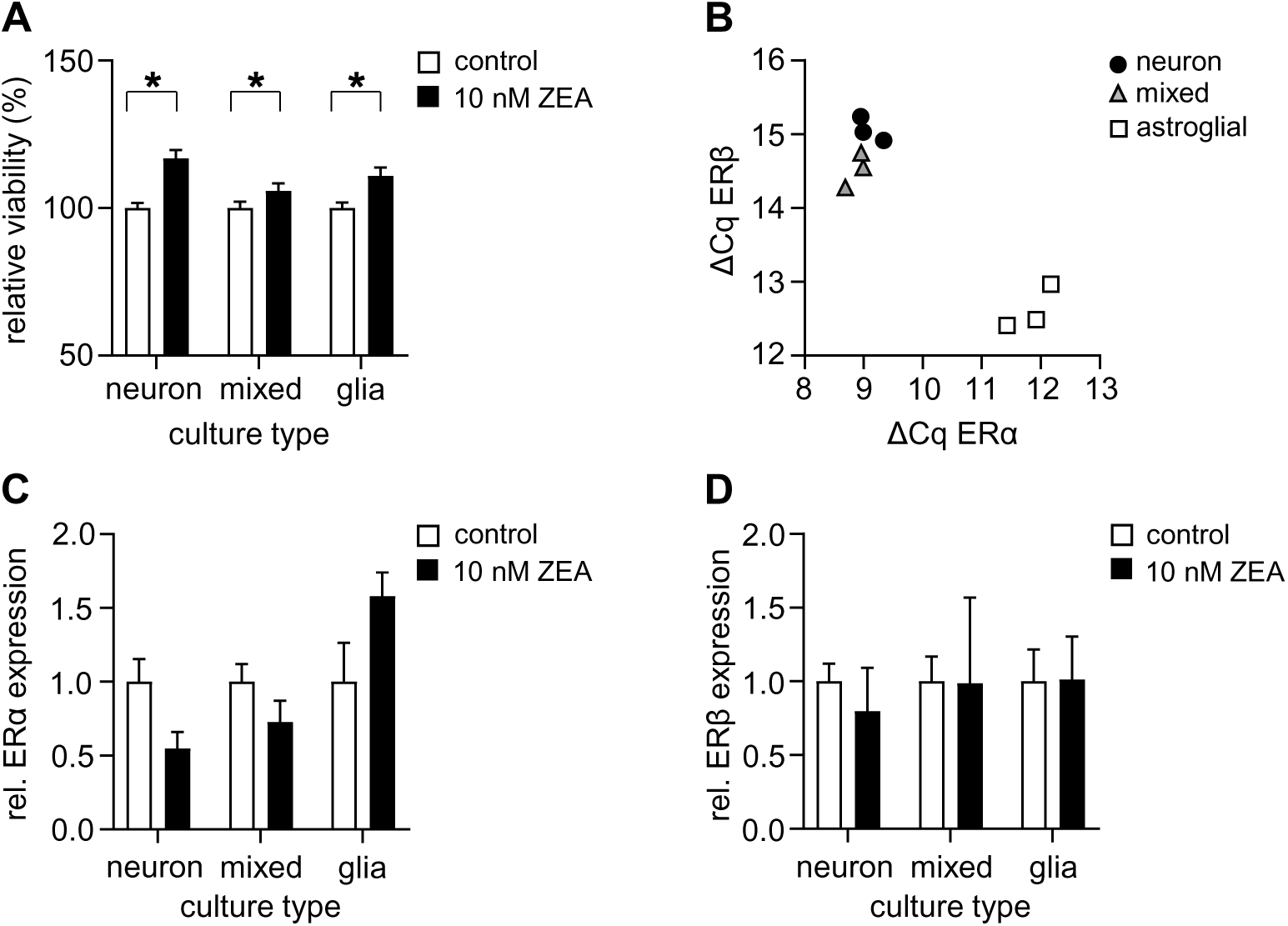
Low ZEA concentration increases cell viability and alters the expression of ERα in a cell type-specific manner. (A) Effects of 10 nM ZEA on the cell viability of mouse primary neuronal, astroglial and mixed cell cultures after 48 hours, as measured by MTT assay. Results are expressed in the percentage of control data, as mean ± s.e.m. n=17-35. * indicates significant differences compared to non-treated data (*:p<0.05). (B) ΔCq values of ERα and ERβ relative expression in the control, non-treated neuron, astroglial and mixed cultures, analyzed by qRT-PCR. Both estrogen receptor types are expressed in the different cell cultures. (B,C) qRT-PCR assay showed that in neuronal and mixed cultures, ZEA administration decreased, while in astroglial cultures, it increased ERα expression compared to the control cells. ERβ expression, on the other hand, was not altered by ZEA in either cell culture type. mRPL13a and GAPDH were used as housekeeping genes. Results are expressed as mean ± s.d. n=3.

As it is known, ZEA, although with higher Kd than estrogen (17β-estradiol, E2; (Nikov et al. 2000) is capable of binding to both ERα and ERβ nuclear estrogen receptors of mammals and induces estrogenic responses at 1-10 nM concentrations. (Shier et al. 2001; Lecomte et al. 2017). Thus, dietary uptake of mycoestrogens may disrupt the natural signaling pathways of E2 (Rai et al. 2019). Based on these former observations, the effect of ZEA on the expression of genes encoding nuclear estrogen receptors was also investigated in all three types of neural cell cultures. First, we examined the transcription rates of the ERα and ERβ genes comparing them to the expression of housekeeping genes (mRPL13a and GAPDH). Analysis of the amplification curves (ΔCq) revealed that both types of estrogen receptors were expressed in both neurons and astroglia cells (Fig. 4B) but their amounts differed between culture types (Fig. 4B). No significant difference in the amount of ERα and ERβ receptors was detected in the astroglial cultures (Fig. 4B). In cultures containing neurons, on the other hand, higher expression of ERα and lower level of ERβ were observed compared to astrocytes (the amplicon lengths were nearly the same, see Suppl. Table 1).

To evaluate the effects of ZEA administration, we quantified the expression of ER mRNAs following 48 hours treatment with 10 nM ZEA, normalized to the corresponding control ΔCq values. According to our data, the expression of ERα was decreased in pure and mixed neuronal cultures and increased in astroglial cultures (Fig. 4C). ERβ expression, on the other hand, was not altered by ZEA treatment in either culture types (Fig. 4D).

As changes in the expression of ERs is generally linked to E2-pathway induction (Liu and Shi 2015), the observed increase in cell viability after ZEA administration suggest metabolic changes in action.

### ZEA does not affect the amount of mitochondria in astrocytes

Estrogens are well known to influence mitochondrial activity under both normal physiological and pathological conditions such as by controlling the transcriptional regulation of mitochondrial proteins, interacting with mitochondrial ER receptors and stabilizing the activity and structural morphological integrity of the mitochondrial respiratory electron-transport chain (Arnold and Beyer 2009). To decide whether the observed increase in cell viability caused by ZEA-treatment is due to a change in mitochondrial function or volume, the extent of the mitochondrial network was estimated in the presence or in the absence of ZEA in astroglia cells (Fig. 5A). To do so, the mitochondrial network was stained with MitoTracker Orange, and its area was normalized to the area of the cytoplasm. Our analyses did not reveal any significant differences in the mitochondrial area compared to the control cultures (Fig. 5B). For further analyses, we also compared the expression of the mitochondrial marker voltage-dependent anionic channel (VDAC1) by qRT-PCR. Relative expression of VDAC1 was compared to the expression of housekeeping genes (mRPL13a and GAPDH). As it is shown on Fig. 5C, ZEA treatment did not substantially alter the expression of the VDAC1 gene. No changes were observed in mixed and astroglial cultures, however, in neuronal cultures, the expression of VDAC1 gene was slightly but not significantly reduced.

**Figure 5.**
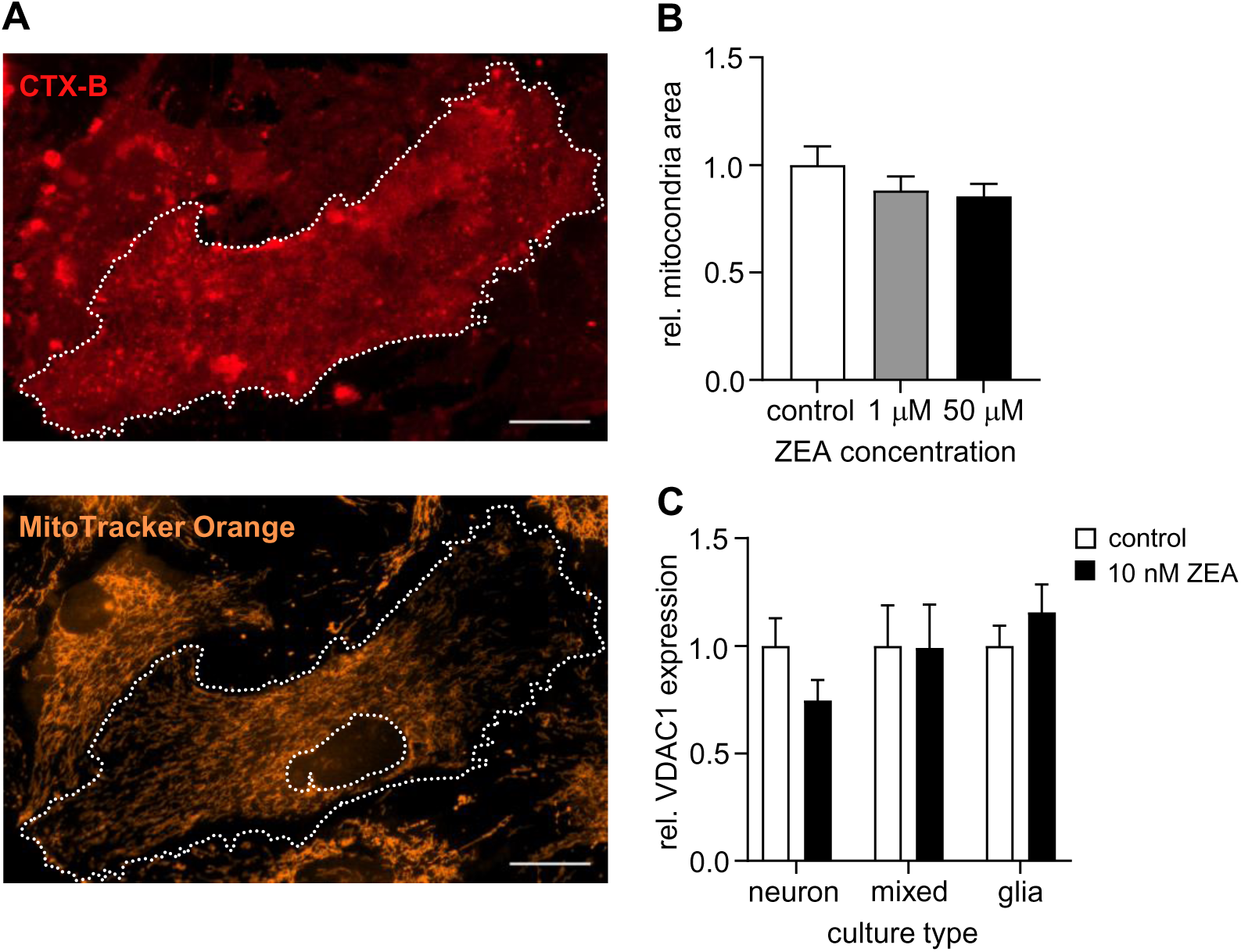
ZEA does not influence the amount of mitochondria but reduces the expression of VDAC1, an ion channel of the outer mitochondrial membrane. (A) CholeratoxinB-Alexa647 was used to visualize the cell membrane, while mitochondria and nuclei were stained by MitoTracker Orange7510 and DAPI, respectively. The relative area of mitochondria was determined as the ratio of mitochondria/cytoplasm areas, using ImageJ software. Scale bars, 20 mm. (B) Effects of 10 nM of ZEA on the relative amount of mitochondria in astroglial cultures. Results are expressed as mean ± s.e.m. (n=12-15). (C) VDAC1 relative expression in ZEA-treated cells compared to the control, non-treated cultures, and analyzed by qRT-PCR. mRPL13a and GAPDH were used as housekeeping genes. Results are expressed as mean ± s.d. n=3.

Taken together, neither immunocytochemical evaluation nor mitochondrial marker gene expression assays were able to detect significant changes in the mitochondrial network in any of the investigated culture types after ZEA treatment. Thus, our results suggest that increased cell viability may be primarily due to other metabolic causes.

### ZEA alters spontaneous neuronal activity in mixed cell culture

It is also known that estrogen can alter the firing activity of neurons (Woolley 2007). To investigate whether the mycoestrogen ZEA has a similar effect on neuronal cultures, multi-electrode arrays (MEAs) electrophysiology was performed to investigate spontaneous electrical activity of neuronal cultures over time (Johnstone et al. 2010). Dissociated neurons were cultivated on top of the electrode surfaces of MEA chips (Fig. 6A). Following a baseline recording, 2nM ZEA was added to the cultures and spontaneous neuronal activity was monitored in a non-invasive way over time, based on changes in local field potentials (LFPs). In short, MEA recordings were performed after 15 minutes and 24 or 48 hours later.

**Figure 6.**
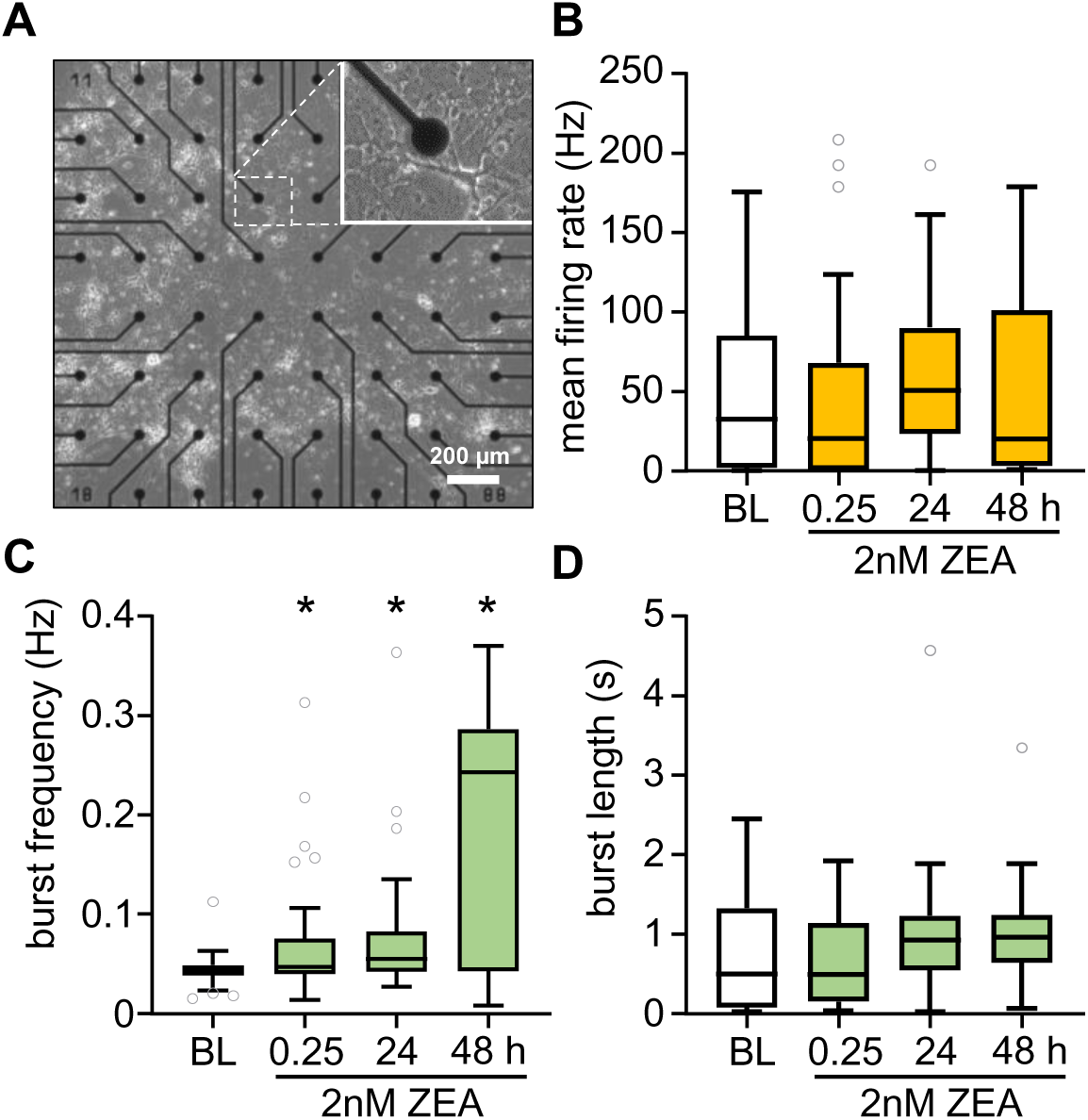
ZEA alters the neuronal activity by affecting firing pattern. (A) Overview of in vitro developed neuronal network, cultured on the surface of a MEA chip. (B-D) Electrophysiological parameters measured by multielectrode array before and after (15 min, 24 h and 48 h) 2 nM ZEA-treatment. Results are represented as box plots (interquartile range and median). *:p<0.05, n=16-40.

Primary neurons grown on MEA chips formed dense neural networks (Fig. 6A) and exhibited robust bursting activity, characterized by high level of the spikes within burst parameter (74.3±5.0%, mean±s.e.m, n=48). ZEA treatment did not change the firing activity, so the mean frequency (Fig. 6B) and the burst length parameters (Fig. 6D) were unaffected upon ZEA treatment. However, the pattern of spontaneous neuronal activity significantly changed as burst frequency values increased (Fig.6C). The effect was already observed with short-term treatments (15 min), which was further enhanced by the long-term presence of ZEA (48 h). The ratio of spikes within bursts was also slightly increased (81.0±6.8%, mean±s.e.m, n=19), but this change was not significant.

These data show that ZEA can influence the spontaneous electrical activity of neuronal networks.

## Discussion

Mycotoxins produced by *Fusarium* spp. molds are worldwide pollutants of food and feed (Marin et al. 2013). Most of them is known to have wide range of biological effects, however their neurotoxic potential and the underlying mechanisms of their toxicity are not so clear. The literature is contradictory regarding their toxic effects in the brain due to the diversity of *in vivo* and cellular models used. In addition, certain species have different tolerance to these toxins (Rotter et al. 1996; Pestka 2007). In this study, we aimed to compare the effect of fumonisin B1, deoxynivalenol and zearalenone on mouse primary cultures containing neurons and astrocytes in different amounts to characterize the primary cellular targets of the different mycotoxins. Moreover, in case of ZEA, we performed detailed analyses to clarify the mode of its action in our *in vitro* system.

### Effects of fumonisin B1 on neural cultures

Previously, several studies showed that FB1 has neuronal effects as it evokes neurotoxicity, caused by oxidative stress or apoptosis (Doi and Uetsuka 2011) and alteration in myelin formation (Monnet-Tschudi et al. 1999). In contrast, Bódi et al. showed recently only a mild increase in neuronal intrinsic excitability *in vitro* as only high concentration (100 μM) of FB1 evoked significant increase in the electrophysiological properties of either hippocampal cell cultures or brain slices. Moreover, changes in neuronal activity and excitability under *in vivo* condition were not detected (Bódi et al. 2020). In addition, FB1 did not cause cell death in astroglia cultures within the 1nM-50μM concentration range used in our experiments, but slightly and significantly increased cell viability. This observation is in line with previous studies. For example, Kwon et al. showed that FB1 (0.5-75 µM) was not cytotoxic after 5 or 10 days of exposure in primary rat astro- and oligodendroglial cultures (Kwon et al. 2000). Similar results were obtained from differentiated primary rat astrocyte cells (Galvano et al. 2002) or in a glioblastoma cell line (Stockmann-Juvala et al. 2004a). However, incubation with 50 µM FB1 for 8 days could cause cell death in primary mouse astrocyte cultures (Osuchowski and Sharma 2005).

In case of neuronal and mixed cultures, the toxin also caused a small (<20%), but significant, dose-independent increase in cell viability at low concentrations (<10 µM). It is consistent with the findings of Harel et al., where primary rat hippocampal cultures did not show signs of cell death despite 48 hours application of FB1 at 10-40 µM concentrations (Harel and Futerman 1993). Other reports also showed non-toxic effects on mixed culture-type even at higher (200 μM) concentrations (Monnet-Tschudi et al. 1999; Domijan and Abramov 2011; Domijan et al. 2012). Notably, FB1 was not toxic to human SH-SY5Y neuroblastoma cells at 200 µM after 24 hours (Domijan and Abramov 2011), however other study reported toxicity over 100 µM after 48 hours (Stockmann-Juvala et al. 2004b).

In our pure neuronal cultures, however, 50 µM FB1 drastically reduced the cell viability. This effect was independent of the duration of treatment (24 vs. 48 hours) and was not observed in the presence of astroglial cells (mixed culture). In a recent study, Domijan et al. reported that besides modifying membranes and induction of oxidative stress, FB1 can inhibit the mitochondrial transport chain I complex and deregulate calcium signaling (Domijan and Abramov 2011). They observed that FB1 administration (0.5-200 μM) significantly increases the level of glutathione (GSH) in astrocytes. As GSH is the major antioxidant in the CNS and astrocytes are the major source of GSH in brain (Dringen and Hirrlinger 2003), the excreted GSH by astrocytes can reduce the oxidative stress in neurons (Bolaños 2016).

Taken together, these results indicate that fumonisin B1 at higher concentrations is more cytotoxic to neurons and the presence of astrocytes can partially compensate the neuronal toxicity.

### Effects of deoxynivalenol on neural cultures

Deoxynivalenol is one of the most abundant Fusarium mycotoxins. In the last two decades, several studies indicated DON contamination in harvested cereals and animal feeds (Tima et al. 2016a, b) and revealed exposures above the tolerable daily intake (TDI: 1μg/ bw kg) to DON in the German and Austrian populations (Warth et al. 2012; Gerding et al. 2014) where the provisional daily intake (PDI) exceeded the TDI in 12% or 33% of the volunteers, respectively. As some effects triggered by DON can occur *in vivo* in the brain at concentrations close to the TDI (Maresca 2013), neurological damage is a risk. Former studies showed that DON can reach neuronal tissues (Prelusky et al. 1990; Pestka et al. 2008) and in concentrations exceeding 10μM, it can reduce the integrity of the BBB (Behrens et al. 2015). It is also known that DON decreases the brain protein levels and the activity of the antioxidant system and induces oxidative stress, apoptosis (Ren et al. 2016) and can cause neuroinflammation (Girardet et al. 2011a). DON has also been reported to have neuroendocrine effects, as well (Rotter et al. 1996).

Based on our results obtained from primary neural cultures, DON acts in a cell-type specific manner. Cell viability was reduced in all three culture types in a concentration- and exposure-time-dependent manner, however, astroglial cells were found to be the most sensitive to DON (cell viability was decreased above 0.2μM concentrations in glial and above 1 μM doses in mixed and neuronal cultures, respectively). Sensitivity of glial cells is consistent with previous work where DON was found to be cytotoxic in rat glia cell cultures *in vitro* within 10-100 µM (Razafimanjato et al. 2011). In this study, DON also reduced the viability of astroglial cells: the IC_50_ dose calculated for the glial cells (31 μM) is comparable with our finding (24 μM). However, in this publication microglial cells were primarily the most sensitive population. Similar effects on the human astroglial cell line, STTG-1 were also demonstrated, but only at higher doses (>25 μM). In addition, DON has also been shown to inhibit the glutamate uptake of astrocytes, which is a key astroglial function (Razafimanjato et al. 2011).

The effect of DON on the viability of neurons obtained from rodents is, to our best knowledge, a new, unpublished scientific result. Recently, Wang et at. presented neuronal data of DON toxicity using piglet hippocampal cells and PC12 cell lines (Wang et al. 2016, 2018). The piglet neurons were already sensitive to DON from ~400nM which, can be explained by the fact that swine is more susceptible to DON than mice or rats (Rotter et al. 1996). Apoptosis and neuroinflammation are the most feasible mechanisms behind the neurotoxicity of DON (Razafimanjato et al. 2011; Wang et al. 2018). According to our data, DON acts in a cell-type specific manner, it is likely that actual tissue concentration of DON determines glial and/or neuronal effects.

### Effects of zearalenone on neural cultures

Little is known about the direct effects of short or continuous exposure to ZEA mycoestrogen on brain function and development. Former publications showed that beside the oxidative- and DNA damage, apoptosis or necrosis caused by ZEA, its main prominent action is through the estrogenic signalization pathways (Rai et al. 2019). Both types of nuclear estrogen receptors, ERα and ERβ, are expressed and play a vital role in neuronal development, and their expression is dynamically regulated during critical periods of cortical development (Belcher 1999). Under physiological conditions, E2 is able to bind to the α-receptor with greater affinity (Kuiper et al. 1997) and E2 induced changes in the expression of estrogen-sensitive genes ultimately lead to regulation of central nervous system development and neuronal differentiation, and affect cell migration, viability and cell death and synaptic plasticity (Beyer 1999).

In this study, expression of both estrogen receptor mRNAs (ERα and ERβ) was detectable by RT-qPCR in all culture types. The expression of ERβ receptors was lower than ERα receptor levels in neuron containing cultures. Similar observations are reported in the literature (Su et al. 2001; Wilson and Westberry 2009; Wilson et al. 2011; Piechota et al. 2017) with the note that the pattern of mRNA expression and protein expression of these receptors is generally correlated (Wilson and Westberry 2009). In the adult brain, E2 has been shown to mostly reduce the expression of ERα mRNA (Liu and Shi 2015). 10 nM ZEA exhibits similar effects in neuronal cultures after 48 hours of exposure. In contrast, in astroglial cultures, ZEA caused a significant increase in ERα expression. This is consistent with Frago et al., who reported that E2 increased the amount of ERα mRNA in astroglial cultures obtained from female rats (Frago et al. 2017). Although in their study, E2 reduced the expression of ERβ in astroglial cells, as well, we did not detect a similar effect of ZEA treatment in mouse astroglial cultures. In mixed cultures, no significant change in one of ER-receptor-mRNA levels was detected after ZEA treatment. Since the sex of the pups used to establish the cell cultures was not determined, further investigations are needed to investigate if ZEA exerts sex-dependent effect on neurons, like E2. It should be noted that in other systems e.g. neuroblastoma cell lines or cerebellar granule cells exhibited opposite effects after ZEA administration (Schreihofer 2005; Jocsak et al. 2016) by strengthening the cell-type specific action of ZEA.

Few studies investigated the direct toxic effects of ZEA on neural cell types (Venkataramana et al. 2014; Kiss et al. 2018; Rai et al. 2019; Agahi et al. 2020a). In our *in vitro* assay, ZEA increased cell viability in all cell culture-types, even at 10 nM. Theoretically, higher cell viability can be achieved by increased proliferation and/or metabolism. Neurons are non-dividing cells excluding the proliferating effect. It is also known that estrogens control the transcriptional regulation of mitochondrial proteins and influence mitochondrial activity under both physiological and pathological conditions via ERβ receptors by e.g. affecting the activity of mitochondrial respiratory electron transport chain, reducing ROS-production or balancing the expression of fusion and fission proteins and changing the structure of mitochondrial network (Arnold and Beyer 2009). According to our results, ZEA dependent changes in the extent of mitochondrial network (area or VDAC1 expression) were not detected. Alternatively, exposure to ZEA can interfere with the oxidoreductive metabolism of mitochondria and may affect the reductive capacity of the cells by increasing SOD and GSH levels (Dong et al. 2007; Tatay et al. 2017; Agahi et al. 2020a). On the other hand, in neuroblastoma cell lines, ZEA induced oxidative stress and reduced cell viability (Venkataramana et al. 2014) only in 1000× higher, presumably not physiological concentrations (Shin et al. 2009b, a).

Importantly, effects of ZEA on neuronal activity and network firing pattern is a new finding. The generally observed burst firing in our cultures are driven by excitatory network episodes due to the dense glutamatergic connections among the cells (Suresh et al. 2016). ZEA has potentiating effect by increasing the frequency of bursts but it does not influence the overall firing rate of the network. This phenomenon resembles the effects of estrogen on neuronal cultures where burst activity has been shown to be more apparent in the presence of estrogen (Brewer et al. 2008).

Our results strengthen the cell type specific effects of *Fusarium* toxins on the nervous system. Data obtained from cell cultures with different neuronal and/or astroglial composition support the importance to identify the cellular targets of *in vivo* mycotoxin exposure.

## List of abbreviations

FB1: Fumonisin B1
DON: deoxynivalenol
ZEA: zearalenone

## Acknowledgments

The authors thank Attila Szűcs for their critical reading and suggestions.

## Funding

This work is supported by grants from the National Research, Development and Innovation Office (PD137855 to N.B., NVKP_16-1-2016-0016, VEKOP-2.3.3-15-2016-00007) and by the National Brain Research Program (2017-1.2.1-NKP-2017-00002).

## Author Contributions

V.S., N.B., K.S. and K.T. wrote the manuscript. V.S. and B.M. performed the cell viability assays and analyses. V.S. and K.T. did the microscopic and qRT-PCR analyses. A.R. measured and analyzed the electrophysiological recordings.

## Conflicts of interest

The authors declare that they have no conflict of interest.

## Declarations

### Availability of data and material

The datasets generated during and/or analyzed during the current study are available from the corresponding author on reasonable request.

### Code availability

Not applicable

### Ethics approval

All experiments complied with local guidelines and regulations for the use of experimental animals (PEI/001/1108-4/2013 and PEI/001/1109-4/2013), in agreement with EU and local legislation.

### Consent to participate

Not applicable

### Consent for publication

Not applicable

## Notes

### Competing Interest Statement

The authors have declared no competing interest.

