## Supplemental Table 1. for "Cell-type specific effects of *Fusarium* mycotoxins on primary neuronal and astroglial cells"

| gene of interest | primer | sequence | product length (bp) |
|---|---|---|---|
| Mus musculus estrogen receptor 1 (alpha) (Esr1) (NM_007956.5) | forward | ATTATGGGGTCTGGTCCTGC | 187 |
| | reverse | TCTTTCCGTATGCCGCCTTT | |
| Mus musculus estrogen receptor 2 (beta) (Esr2) (NM_207707.1) | forward | CATTACGGTGTCTGGTCCTGT | 189 |
| | reverse | TTCTCTCCTGGATCCACACTTG | |
| Mus musculus ribosomal protein L13A (Rpl13a) (NM_207707.1) | forward | AGGGGCAGGTTCTGGTATTG | 191 |
| | reverse | GGGGTTGGTATTCATCCGCT | |
| Mus musculus voltage-dependent anion channel 1 (Vdac1) (NM_001362693.1) | forward | GGCTACGGCTTTGGCTTAAT | 199 |
| | reverse | CAGTGATCTCAGTGCCCAGG | |
| Mus musculus GAPDH (NM_008084.3) | forward | TGGTGAAGGTCGGTGTGAAC | 106 |
| | reverse | ATGAAGGGGTCGTTGATGGC | |

***Supp. table 1.*** Primer sets used in qRT-PCR analysis. The designed primer pairs recognize all transcriptional variants.
